# Single-cell dissection of a rare human prostate basal cell carcinoma

**DOI:** 10.1101/818260

**Authors:** Xianbin Su, Qi Long, Juanjie Bo, Yi Shi, Li-Nan Zhao, Yingxin Lin, Qing Luo, Shila Ghazanfar, Chao Zhang, Qiang Liu, Lan Wang, Kun-Yan He, Jian He, Xiao-Fang Cui, Jean Y. H. Yang, Ze-Guang Han, Jian-Jun Sha, Guoliang Yang

## Abstract

As a rare subtype of prostate carcinoma, basal cell carcinoma (BCC) has not been studied extensively and thus lacks systematic molecular characterization. Here we applied single-cell genomic amplification and RNA-Seq to a specimen of human prostate BCC (CK34βE12^+^/P63^+^/PAP^−^/PSA^−^). The mutational landscape was obtained via whole exome sequencing of the amplification mixture of 49 single cells, and the 5 putative driver genes mutated are *CASC5*, *NUTM1*, *PTPRC*, *KMT2C* and *TBX3*. The top 3 nucleotide substitutions are C>T, T>C and C>A, similar to common prostate cancer. The distribution of the variant allele frequency values indicated these single cells are from the same tumor clone. The transcriptomes of 69 single cells were obtained, and they were clustered into tumor, stromal and immune cells based on their global transcriptomic profiles. The tumor cells specifically express basal cell markers like *KRT5*, *KRT14* and *KRT23*, and epithelial markers *EPCAM*, *CDH1* and *CD24*. The transcription factor (TF) co-variance network analysis showed that the BCC tumor cells have distinct regulatory networks. By comparison with current prostate cancer datasets, we found that some of the bulk samples exhibit basal-cell signatures. Interestingly, at single-cell resolution the gene expression patterns of prostate BCC tumor cells show uniqueness compared with that of common prostate cancer-derived circulating tumor cells. This study, for the first time, discloses the comprehensive mutational and transcriptomic landscapes of prostate BCC, which lays a foundation for the understanding of its tumorigenesis mechanism and provides new insights into prostate cancers in general.

## Introduction

Prostate cancer is a heterogeneous disease with complex subtypes and different cell origins [1–3], and basal cell carcinoma (BCC) is an extremely rare histological subtype comprising < 0.01% of prostate cancer [4]. Despite being largely regarded as pursuing an indolent clinical course, aggressive behaviors like recurrence or metastasis have been observed and deaths also reported, highlighting the complex and yet poorly understood mechanism [5–9]. There have been extensive reports on the genomic features of common prostate cancer [10–17], and the molecular features of normal human prostate basal cells, prostate basal cell hyperplasia, and basal populations from human prostate cancer have also been reported [18–20], but the molecular profile of prostate BCC is still lacking. The mutational and transcriptomic features of this tumor sub-type will thus be valuable for understanding of its tumorigenesis mechanism and development of future clinical treatments.

Single-cell sequencing has evolved to be a powerful tool that provides unprecedented resolution of clinical specimens especially with limited amounts, and marker-free decomposition of the constituent cell types based on single-cell transcriptional profiles allows precise dissection of complex systems such as various tissues or tumors [21–23]. Here we applied single-cell genomic amplification to a human specimen of prostate BCC and used the mixture of 49 single cells to provide mutational landscape of prostate BCC via whole exome sequencing. We then used single-cell RNA-Seq to obtain the transcriptomes of 69 cells from the same specimen. Three types of cells, namely tumor, stromal and immune cells, were identified with their molecular features revealed at single-cell resolution, providing a useful resource for not only this prostate cancer subtype but prostate cancer in general.

## Results

### Histological and immunohistochemical features of human prostate BCC

The tumor exhibited a widespread infiltrative growth pattern under microscopic examination, and individual cells had large pleomorphic nuclei and scant cytoplasm (Fig. 1). The tumor was negative for two known prostatic markers, prostatic acid phosphatase (PAP) and prostate-specific antigen (PSA), but strongly expressed CK34βE12 and P63, suggestive of basal cell origin (Fig. 1). Based on the cellular morphology and immunohistochemical staining, we diagnosed it as prostate BCC. The patient with BCC was resistant to chemotherapy but sensitive to radiotherapy, and followed-up for 32 months without recurrence or metastasis (Supporting Information, Fig. S1).

**Fig. 1.**
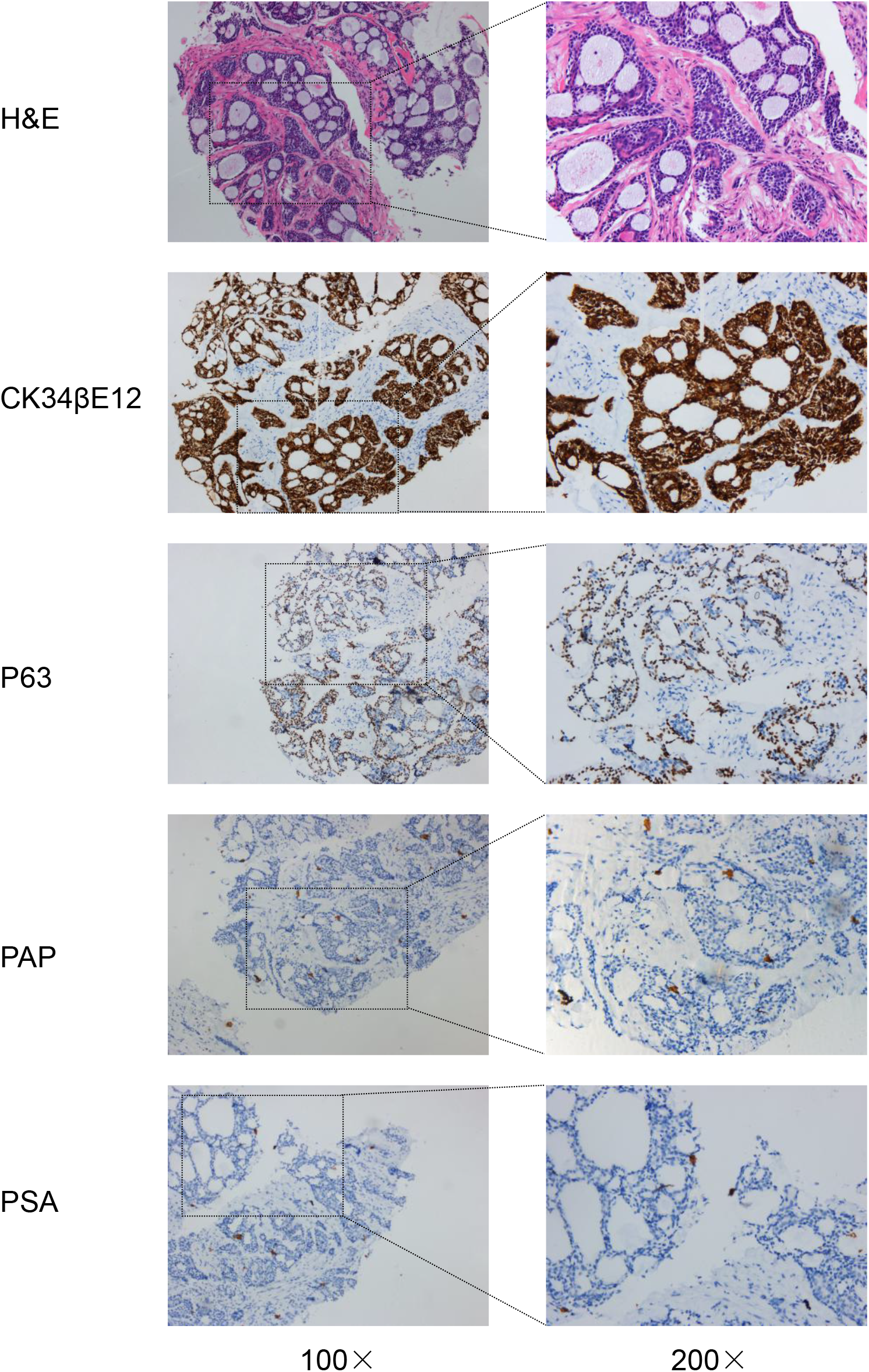
Histological and immunohistochemical staining of the prostate BCC. H&E staining and the expressions of CK34βE12, P63, PAP and PSA are shown.

### Whole exome sequencing reveals the mutational landscape of prostate BCC

As genetic variations are generally believed to be the causes of tumors, here we tried to reveal the mutational features of prostate BCC. Due to the limited amount of tumor specimens collected, after dissociation of the specimens into single-cell suspension, we first conducted single-cell whole genome multiple displacement amplification (MDA) on a microfluidic-chip. We then used the mixture of amplified products from 49 single cells for whole exome sequencing (Fig. 2A). A total of 91 functional exonic mutations were obtained, and 5 of the mutated genes, *CASC5*, *NUTM1*, *PTPRC*, *KMT2C* and *TBX3* are putative driver genes catalogued in the COSMIC Cancer Gene Census (Supporting Information, Fig. S2) [24]. Mutations in DNA repair genes are related to poor outcomes of prostate cancers [12, 25], but few are mutated in this prostate BCC sample such as *BRCA2*, consistent with good survival of this patient.

**Fig. 2.**
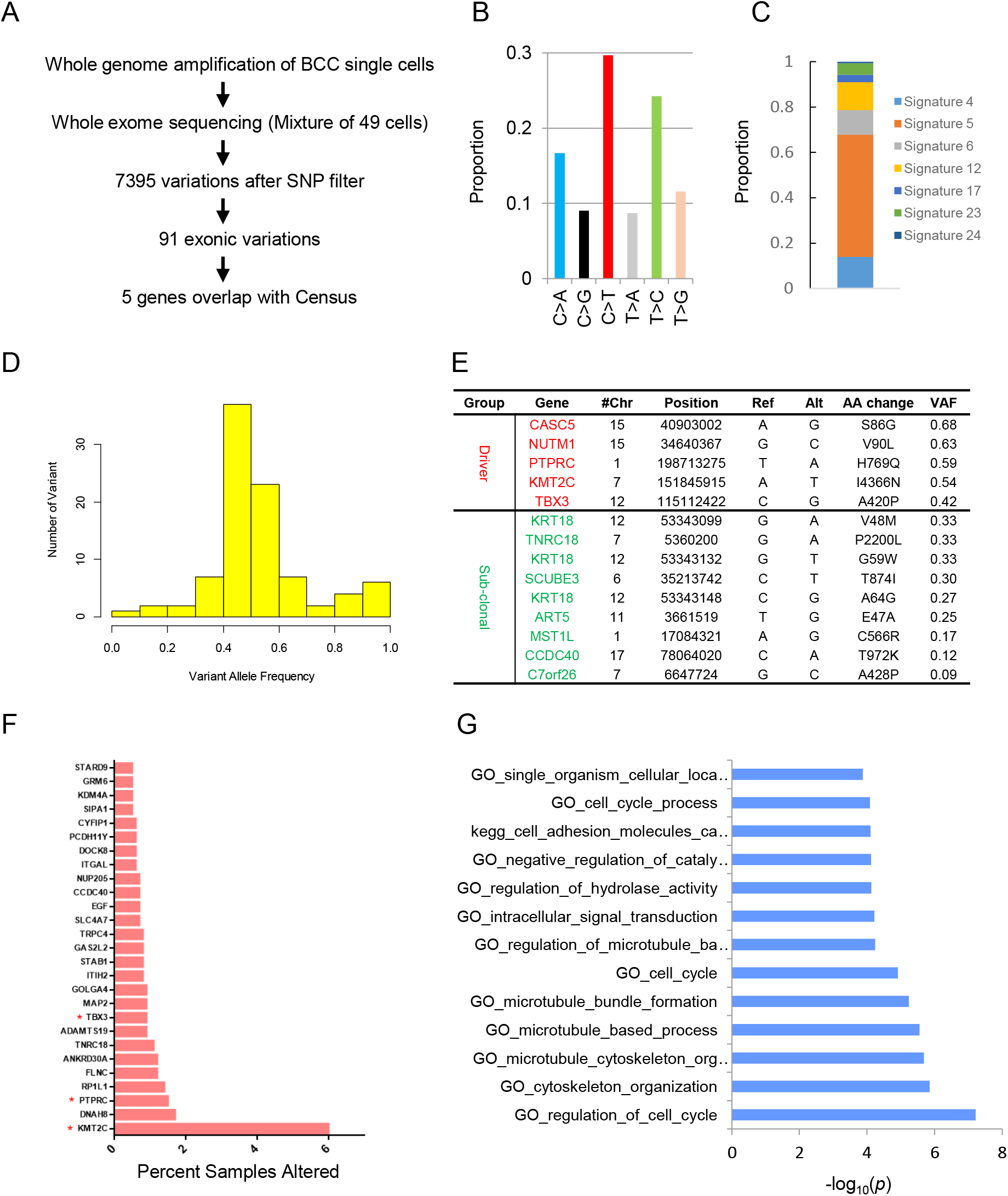
Whole exome sequencing reveals the mutational landscape of prostate BCC. General description of the exome sequencing of mixtures from genomic amplification products of prostate BCC single cells and numbers of genetic variations. Mutational spectra of the prostate BCC based on the variants after SNP filter. (C) Contributions of COSMIC signatures to the prostate BCC mutational spectra. (D) Distribution of the variant allele frequency (VAF) of the exonic mutations of prostate BCC. (E) Summary of the driver (overlapped with The Cancer Gene Census) and putative sub-clonal (VAF lower than 0.35) mutations of prostate BCC. (F) Percent of samples mutated in the cBioPortal prostate cancer datasets with those above 1% shown for the exonic mutated genes in prostate BCC. The red stars indicate Census driver genes. (G) The items enriched with the exonic mutated genes in prostate BCC via GSEA/MSigDB canonical pathway enrichment analysis.

The top 3 nucleotide substitutions are C>T, T>C and C>A, which are exactly the same top 3 substitutions in recently reported localized, non-indolent prostate tumors (Fig. 2B) [13]. This suggests the etiology of prostate BCC is likely the same as common prostate tumors. The major composite COSMIC signatures are Signature 4, 5, 6 and 12 (Fig. 2C). While the etiologies of Signature 5 and 12 are still unknown, Signature 4 and 6 are associated with smoking and defective DNA mismatch repair, respectively [26].

By checking the distribution of the variant allele frequency (VAF) of the exonic mutations of prostate BCC, it is clear that most of the variations have VAF values of around 0.5, thus supporting that most of the single cells share a similar mutational background (Fig. 2D). The VAF values of the 5 driver mutations are also approximately 0.5 (Fig. 2E), indicating these single cells are probably originated from the same ancestor and form the major tumor clone. However, there are still some variations with lower VAF values which suggest sub-clonal mutations may have existed in some cells that are acquired during later diversified evolution of the tumor.

To understand whether the prostate BCC is showing similarities to other reported prostate cancer cases, we checked the mutational frequency of the prostate BCC mutated genes in prostate cancer samples collected in cBioPortal [27]. For the 88 genes enquired, only 10 genes are not mutated, and 27 genes are mutated in more than 0.5% of the ~4400 prostate cancer samples (Fig. 2F). Interestingly, 3 of the driver genes from this study, *KMT2C*, *PTPRC*, and *TBX3* are also among the top mutated genes, especially *KMT2C* with mutation rate of ~6%. GSEA/MSigDB enrichment analysis [28] of the mutated genes shows that the top enriched terms are related to cell cycle, cytoskeleton and others (Fig. 2G), consistent with its basal cell origin.

### Single-cell RNA-Seq reveals the transcriptomic features of prostate BCC

To understand the transcriptional profiles of prostate BCC, we then randomly selected 69 single cells for single-cell RNA-Seq (Fig. 3A). With a median unique mapping rate of 58.4% and 1.08 million mapped reads, we obtained a median of ~2800 genes per cell (Supporting Information, Fig. S3A). We used ERCC spike-ins as positive controls, and the high correlation efficiency between single-cells based on the 92 spike-ins confirmed the high quality of the data (Supporting Information, Fig. S3B-D). After filtering 5 outliers, the remaining 64 cells were classified into three clusters based on their global transcriptomic profiles (Fig. 3B-C). GSEA/MSigDB enrichment analysis identified them as tumor cells (51 cells), stromal cells (7 cells) and immune cells (6 cells) (Fig. 3D).

**Fig. 3.**
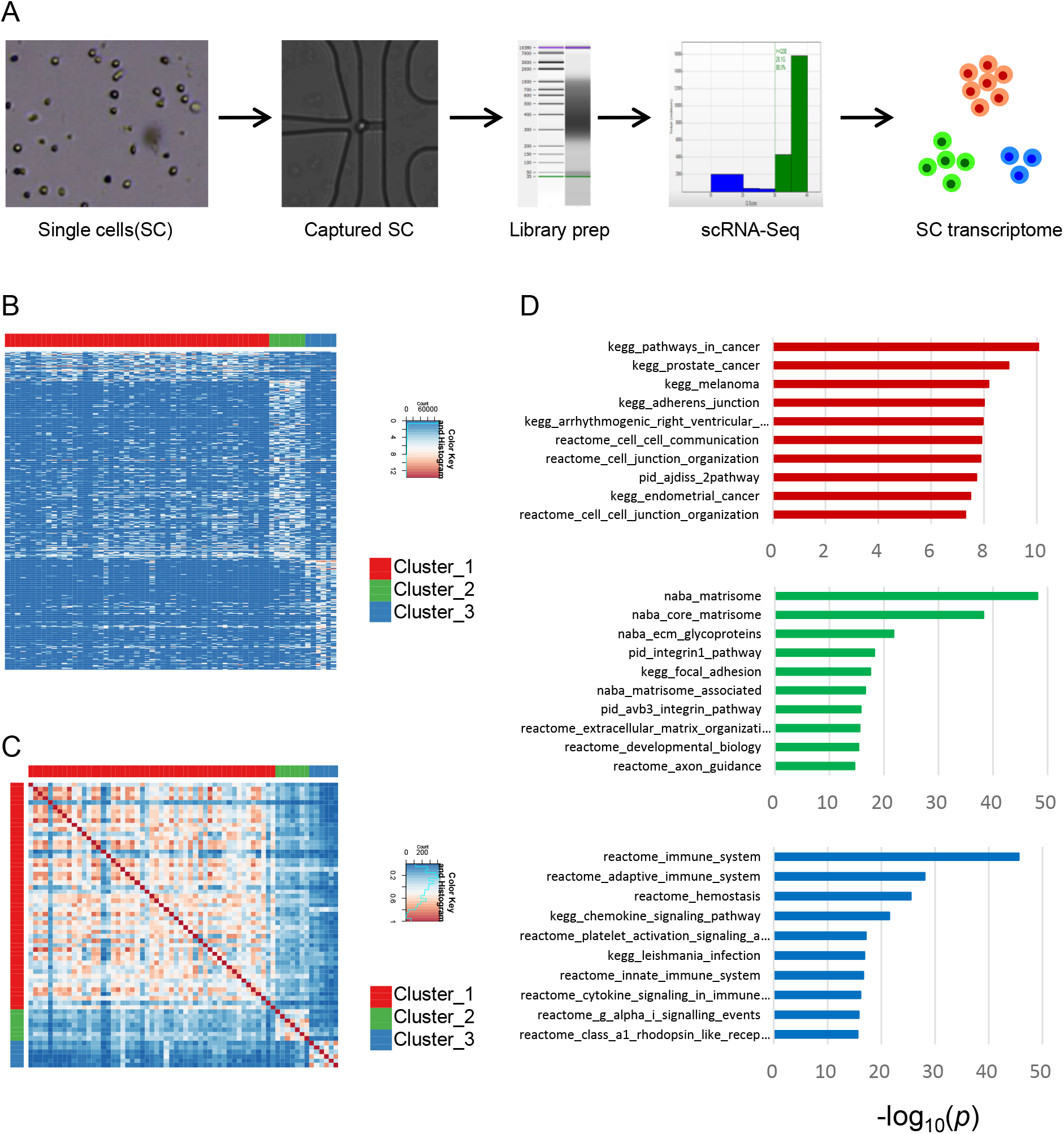
Single-cell RNA-Seq reveals the constituent cell types in the prostate BCC. (A) Experimental workflow. (B) Heat-map showing the single-cell gene-expression patterns of the three clusters identified. (C) Heat-map showing the Pearson correlation coefficients between single-cells. (D) The top 10 items enriched with the gene sets specifically expressed in each cell cluster via GSEA/MSigDB canonical pathway enrichment analysis.

The three clusters of cells can also be clearly separated in tSNE plot based on the top genes expressed in each cluster (Fig. 4A). The tumor cells specifically expressed genes encoding cytokeratins including *KRT5*, *KRT14* and *KRT23*, which are reliable basal cell markers (Fig. 4B). Interestingly, epithelial markers *EPCAM*, *CDH1* and *CD24* are also highly expressed in tumor cells. Stromal cells specifically express *COL1A2*, *THY1* and *BGN*, and immune cells specifically express monocytic marker *CD14*, *CXCL8* and myeloid cell nuclear differentiation antigen *MNDA* (Fig. 4B), suggestive of possible innate immune cell infiltration. Immunohistochemical staining confirmed the expression of CK14 (KRT14) at protein level, further proving the reliability of the sequencing data (Fig. 4C).

**Fig. 4.**
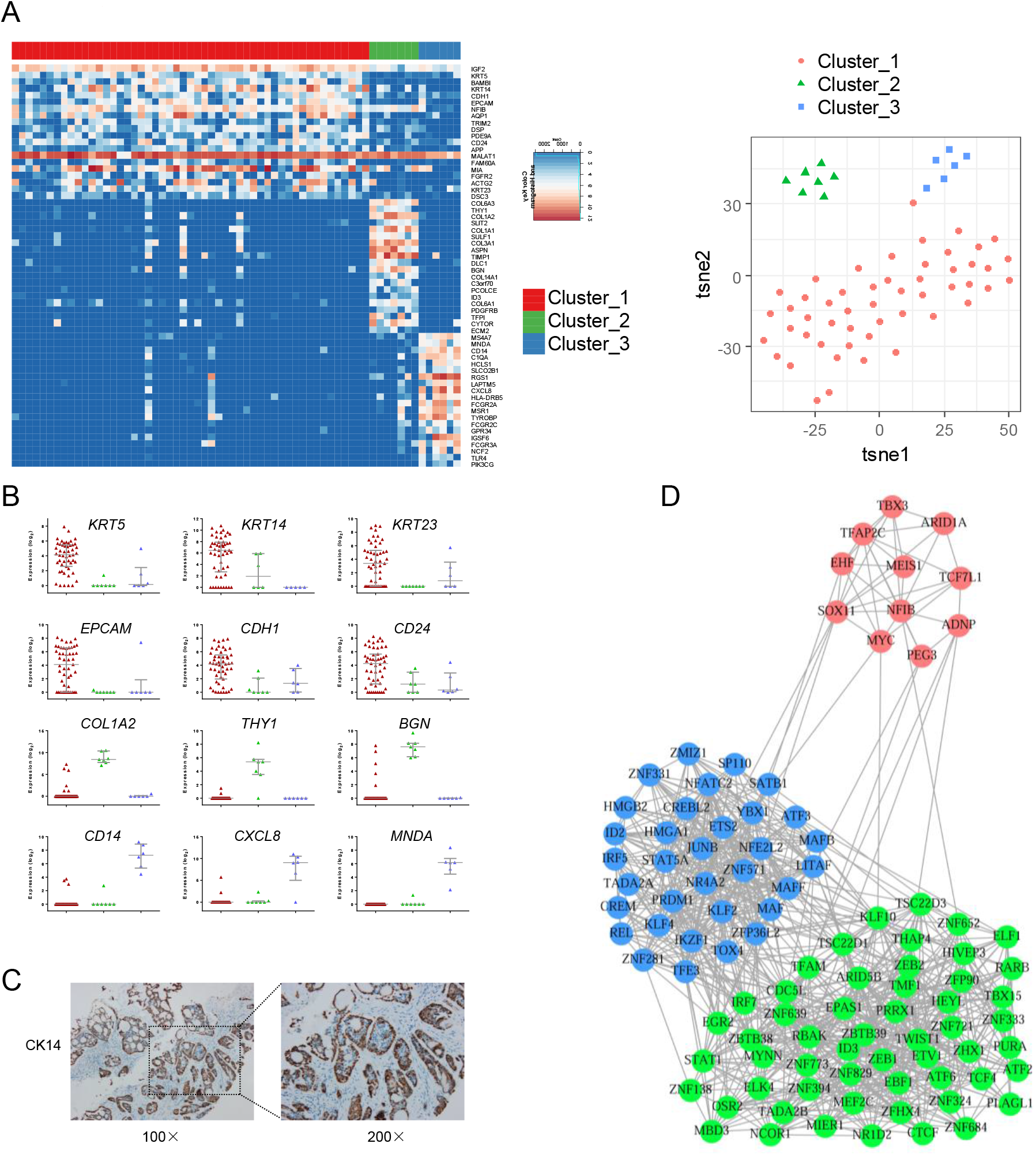
Single-cell RNA-Seq reveals the transcriptomic features of prostate BCC. (A) Heat-map showing the expression patterns of top 20 genes specifically and highly expressed in each cluster, and t-SNE plot of the three clusters. (B) The expression of selected markers with the median values and the first and third quartiles shown. (C) Immunohistochemical staining of the specimen with CK14 antibody. (D) TFs covariance networks of the single-cell data. Each node represents a TF, and each edge represents correlation between two TFs. The TFs specifically expressed in each cluster are colored differently.

Transcription factors (TFs) covariance network analysis showed that the three clusters demonstrate different transcriptional regulatory status (Fig. 4D). Significantly, *MYC*, *ARID1A*, *TBX3* and *SOX11* are highly expressed in tumor cells, where, besides the well-known *MYC* that is recently reported to be over-expressed in human prostatic basal cells [19], the oncogenic role of *ARID1A*, a key component of chromatin remodeling complex SWI/SNF, has been recently demonstrated in liver cancer [29], the development-related gene *SOX11* was reported to be elevated in basal-like breast cancer [30], and *TBX3* plays important roles in development, differentiation and tumorigenesis [31]. The TFs co-variance network provides clues to future dissection of the regulatory mechanism of BCC.

### Single-cell analysis of prostate BCC provides new insights to prostate cancer

We then checked the expression of prostate BCC tumor cell specific genes in previously reported luminal and basal cells sorted from normal and cancerous prostates by flow cytometry [18]. The prostate basal cells exhibit higher expression levels of *KRT5*, *KRT14* and *KRT23* compared with luminal cells, similarly as our prostate BCC tumor cells, further confirming their basal cell origin (Fig. 5A). Interestingly, the genes highly expressed in prostate BCC tumor cells fall into two groups based on their expression correlation patterns, with one group including *KRT5*, *KRT14*, *KRT23, DSC3, FGFR2*, etc., while the other group including *EPCAM*, *CDH1*, *CD24*, etc. (Fig. 5A). It seems the former group includes the basal cell markers while the latter group includes genes expressed in common prostate epithelial cells.

**Fig. 5.**
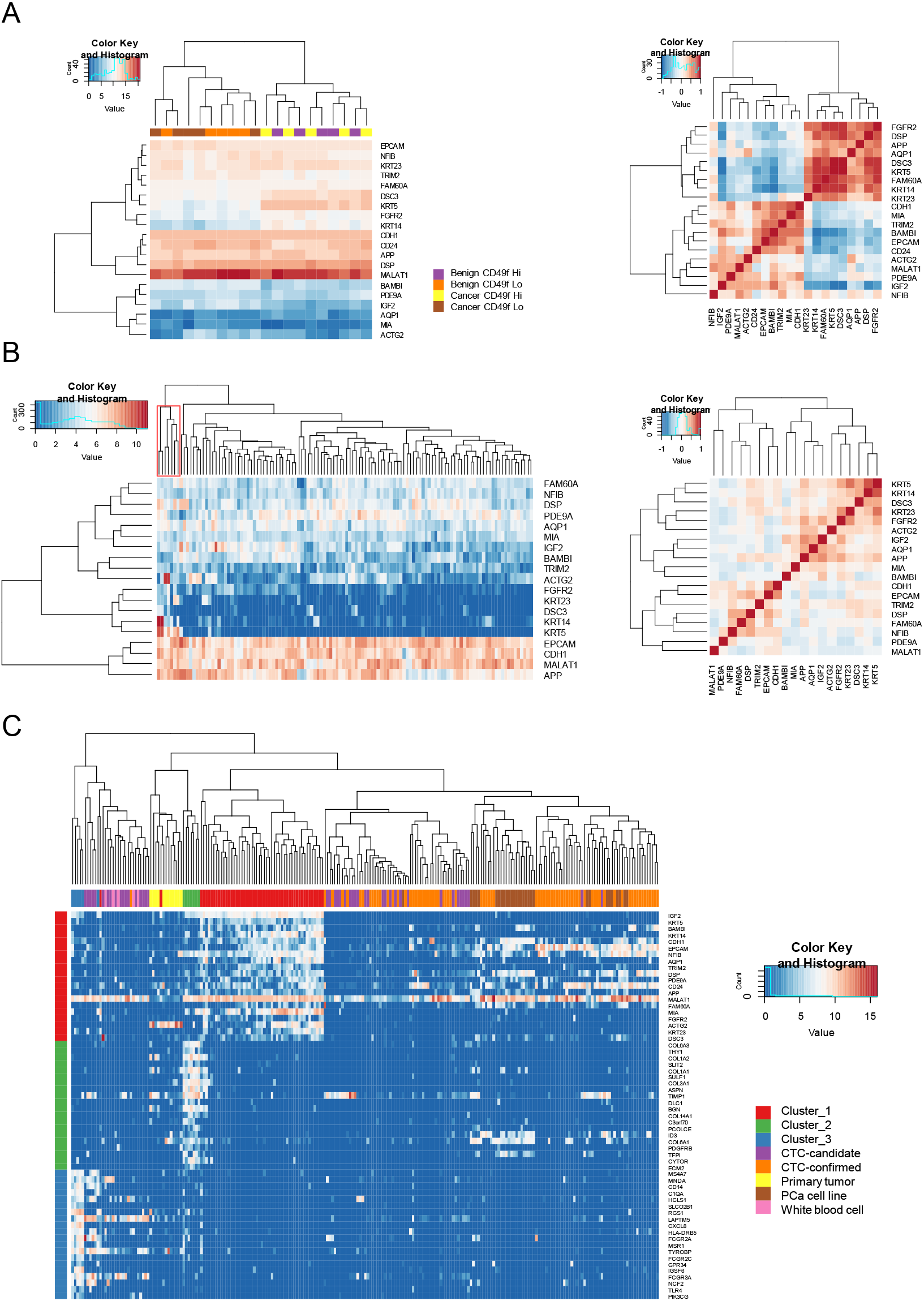
Single-cell analysis of prostate BCC provide new insights to prostate cancer. (A) (left) Heat-map showing the clustering of FACS sorted luminal (CD49f Lo) and basal (CD49f Hi) cell populations from human benign and cancerous prostates based on the genes highly expressed in prostate BCC tumor cells; (right) Heat-map showing the Pearson correlation coefficients between the genes. (Data from Smith *et al*. PNAS 2015; 112: E6544-E6552). (B) Analysis of prostate cancer samples in similar approach as (A). (Data from Robinson *et al*. Cell 2015; 161: 1215–1228). (C) Comparison of the gene expression profiles of single cells from Cluster 1~3 of prostate BCC with human prostate CTCs (CTC-candidate and CTC-confirmed), bulk primary tumors, single cells from prostate cancer cell lines (Pca cell line) and white blood cells (Data from Miyamoto *et al*. *Science* 2015; 349: 1351-1356).

We further checked the expression patterns of BCC tumor cell specific genes in other bulk samples of human prostate cancers [10, 11, 32]. It is interesting to identify a small group of samples exhibiting basal cell gene expression features in two separate prostate cancer datasets (shaded branches in the dendrograms of Fig. 5B and Supporting Information, Fig. S4A). Likely explanations are that these tumor samples were derived from BCC patients or the samples contained basal cells. For TCGA PRAD dataset, the expressions of basal markers are detected in most samples, possibility due to the high number of genes detected by deep sequencing (Supporting Information, Fig. S4B). The split of the genes highly expressed in BCC tumor cells into 2 groups is further confirmed by these three datasets, with *KRT5*, *KRT14*, *KRT23* clustering together with *DSC3, FGFR2* and also possibly *ACTG2.* Our single-cell analysis of prostate BCC reveals the existence of basal cell features in current bulk samples of prostate cancer.

We then compared our results with one single-cell resolution dataset of prostate cancer [33]. Compared with circulating tumor cells (CTCs) from prostate cancer patients, the prostate BCC tumor cells exhibit distinct gene expression features with high levels of *IGF2*, *KRT5*, *KRT14*, *KRT23*, *AQP1*, *MIA*, *FGFR2* and *ACTG2*, although they all expressed *EPCAM*, *CDH1* and *CD24* (Fig. 5C). The results demonstrated the unique gene expression patterns of the prostate BCC tumor cells, which exhibit the molecular features of basal cells with activated genes like *IGF2* and *FGFR2*, consistent with report on the regulation of pluripotency by IGF and FGF pathways [34]. Another interesting finding is that the BCC immune cells clustered together with some single cells annotated as CTC-candidate, which were likely to be contaminating leukocytes to the CTCs (Fig. 5C). The single-cell level data also support the above mentioned split of the genes highly expressed in prostate BCC tumor cells, with one group representing genuine BCC markers while the other genes commonly expressed in prostate cancer.

### Detection of exonic mutations in single-cell RNA-Seq reads

As our mutational data and single-cell RNA-Seq data were generated from the same prostate BCC specimen, we checked whether the exonic mutations could be detected in RNA-Seq reads. Interestingly, we indeed found mutations in RNA-Seq reads from some single cells for *GABPB2*, *ASTE1*, *COPS3* and *BEX2* (Supporting Information, Fig. S5). The variations are only detected in some of the single cells, which are likely because the mutated sites are not always covered by the relatively shallow single-cell RNA-Seq. Another phenomenon is that single cells in RNA-Seq analysis only contained either reference or altered variant, which is probably caused by allelic specific expression or allele dropout in RNA-Seq. The consistency of scRNA-Seq data and exome-seq data further suggested the reliability of our prostate BCC data.

## Discussion

As a rare malignancy, there has been little research into the molecular characteristics of prostate BCC and no consensus on its treatment. Mostly, such data exist as sporadic reports on immunohistochemical expression in a small number of cases [8, 35–37]. This study represents the first genomic and transcriptomic profiling of this prostate cancer subtype, and the single-cell resolution transcriptomic data is especially valuable for understanding of not only this subtype but prostate cancer in general.

The molecular features of normal human prostate basal cells [19], prostate basal cell hyperplasia [20], and human prostate cancer derived basal populations [18] have been reported recently, and the single-cell transcriptomic profiles of prostate BCC is a good complement to these data for comparison studies and better understanding of prostate cancer cell origin. Most of the current genomics data for prostate cancer are generated using bulk samples [10, 11, 13], and the single-cell transcriptional profiles of prostate BCC provide an opportunity for deeper utilization of these data. For example, analysis using genes highly expressed in prostate BCC reveals the existence of basal cell features in some bulk samples of prostate cancer, which is consistent with recent classification of prostate cancer samples into basal-like and luminal-like subtypes using the PAM50 classifier [1, 38]. The single-cell resolution profiles of constitutional cell types from prostate BCC provide further advantage for future de-convolution of bulk samples [39], which will maximize the values of current genomic data of prostate cancer.

The prostate BCC patient is still surviving now, indicating this case is an indolent malignancy. Our genomics and transcriptomic analysis of this chemotherapy-resistant but radiotherapy-sensitive prostate BCC provides a useful resource for future dissection of the molecular mechanisms related to the radiotherapy responsiveness and treatment guidance. Mutations that disrupt the function of DNA damage repair genes have been shown to be associated with the aggressive clinical behavior of localized prostate cancer and with cancer-specific mortality [12, 40–42], and it has also been reported that men with metastatic prostate cancer and DNA-repair gene mutations have sustained responses to poly-ADP ribose polymerase (PARP) inhibitors and platinum-based chemotherapy [43, 44]. This study showed that the prostate BCC patient with few DNA-repair gene mutations was resistant to chemotherapy, suggesting that we may directly adopt radiotherapy rather than chemotherapy for such conditions. The results indicated that precision treatment of BCC could be more than just a histopathological triviality but be based on association of tumor phenotypes with molecular features.

In summary, single-cell analysis revealed the genomic and transcriptomic landscapes of the rare prostate BCC for the first time. This provides clues for elucidation of the tumorigenesis mechanism, further discovery of biomarkers and therapeutic targets, and better understanding of not only prostate BCC but prostate cancer in general.

## Methods

### Clinical specimens

This study was approved by the Ethnical Review Board of Renji Hospital, School of Medicine, Shanghai Jiao Tong University. Specimens were obtained by prostatic needle biopsy from a 55-year-old man with pelvic pain and irritative urinary symptoms. The specimens underwent pathological evaluation. After diagnosis as prostate BCC, the patient was treated with chemotherapy and radiotherapy and then followed-up for 32 months without evidence of local recurrence or distant metastases (Supporting Information, Fig. S1). The specimens were mechanically dissociated and digested with collagenase IV and DNase I, and the single-cell suspensions were used for both single-cell genome amplification and single-cell RNA-Seq.

### Immunohistochemical staining

Immunohistochemical staining was done as previously described [45], and we used the following primary antibodies: P63 (abcam, ab124762), CK34βE12 (Dako, M0630), CK14 (abcam, ab7800), PAP (Dako, M0792), PSA (Dako, A056201). We used biotinylated universal link antibody as the secondary antibody.

### Single-cell genome amplification and whole exome sequencing

Single-cell capture, lysis, reverse-transcription, and whole genome MDA were done in a microfluidic-based C1 DNA-Seq IFC (10~17 μm, Fluidigm) according to its protocol using illustra GenomiPhi V2 DNA Amplification Kit (GE Healthcare, 25660031). The amplified products from 49 single cells were mixed for exonic region capture and library preparation using Agilent SureSelect Human All Exon v7 Kit (Agilent, 5191-4005). The library was sequenced using illumina HiSeq platform with 2 × 150 bp sequencing mode. We used GATK [46] for variant calling and ANNOVAR [47] for functional annotation of the mutations. As there is no para-tumor tissue for this patient, we filtered SNPs using dbSNP141 [48] and 1,000 Genomes Project (v3) database [49]. We used MutationalPatterns [50] to decipher the mutational signature composition. For genes mutated in this prostate BCC sample, we checked their mutational frequencies in prostate cancer samples collected in cBioPortal [27]. GSEA/MSigDB enrichment analysis was also conducted for the genes mutated in this prostate BCC.

### Single-cell RNA-Seq and data analysis

Single-cell RNA-Seq experiment and analysis were conducted as previously described [45]. Single-cell capture, lysis, reverse-transcription, and cDNA amplification were done in a microfluidic-based C1 RNA-Seq IFC (10~17 μm, Fluidigm). After library preparation with Nextera XT Kit and Index Kit (illumina), the single-cell libraries were pooled and sequenced using NextSeq 500 (illumina) with 2 × 76 bp sequencing mode. After generation of FPKM data, SC3 [51] was used for identification of cell outliers and single-cell clustering. The genes differently expressed between clusters were then identified via ANOVA analysis (*p* < 0.05) using SINGuLAR™ (Fluidigm). GSEA/MSigDB enrichment analysis was conducted for the genes specifically expressed in each cluster to facilitate cell type identification. Transcription factors (TFs) covariance network analysis was conducted to reveal the relationship of the TFs specifically expressed in each cell cluster as previously described [45, 52]. Gene expression files were retrieved from five other prostate cancer studies [10, 11, 18, 32, 33] for comparison studies.

## Supporting information

Figure S1. Treatment details of prostate BCC patient. Magnetic resonance imaging (MRI) showed a large, irregular tumor mass (arrows)

## Abbreviations

BCC: basal cell carcinoma
CTCs: circulating tumor cells
MDA: multiple displacement amplification
PAP: prostatic acid phosphatase
PSA: prostate-specific antigen
TF: transcription factor
VAF: variant allele frequency

## Acknowledgement

This work is supported in part by the National Natural Science Foundation of China (81802806, 81472621, 81402329 and 81902561), National Program on Key Research Project of China (2016YFC0902701, Precision Medicine), Shanghai Health Bureau (20164Y0124), Medical and Engineering Crossover Fund of SJTU (YG2016QN71, YG2017MS67), University of Sydney – Shanghai Jiao Tong University Joint Research Alliance (USYD-SJTU JRA) grant, funding from Key Laboratory of Systems Biomedicine (Ministry of Education) (KLSB2017QN-03), and Incubating Program for Clinical Research and Innovation of Renji Hospital SJTU School of Medicine (PYII-17-010).

## Authors’ contributions

XS and GY designed the study. XS, JB, LZ, CZ, QL3, LW, KH, JH, XC, WX and GY carried out experiments. XS, QL1, YS, YL, QL2, SG and JY analyzed data. XS, GY and ZH drafted the manuscript. GY, JS and ZH supervised the study. All authors read and approved final version of the manuscript.

