## Supplementary material for "Single-cell dissection of a rare human prostate basal cell carcinoma": Figure S1. Treatment details of prostate BCC patient. Magnetic resonance imaging (MRI) showed a large, irregular tumor mass (arrows)

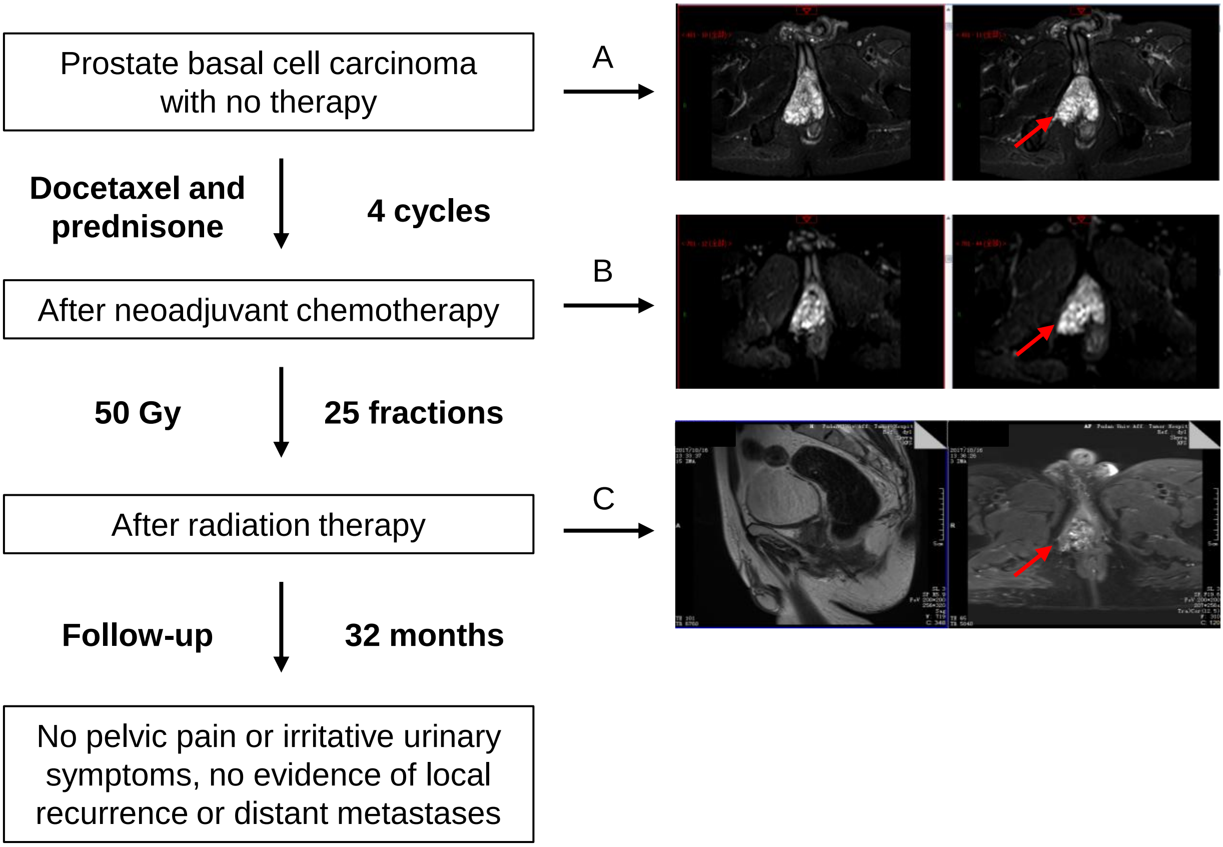


**Figure S1**. Treatment details of prostate BCC patient. Magnetic resonance imaging (MRI) showed a large, irregular tumor mass (arrows): (A) Before treatment; (B) After chemotherapy, there was no significant change in the size of the tumor; (C) After radiotherapy, the tumor shrank.

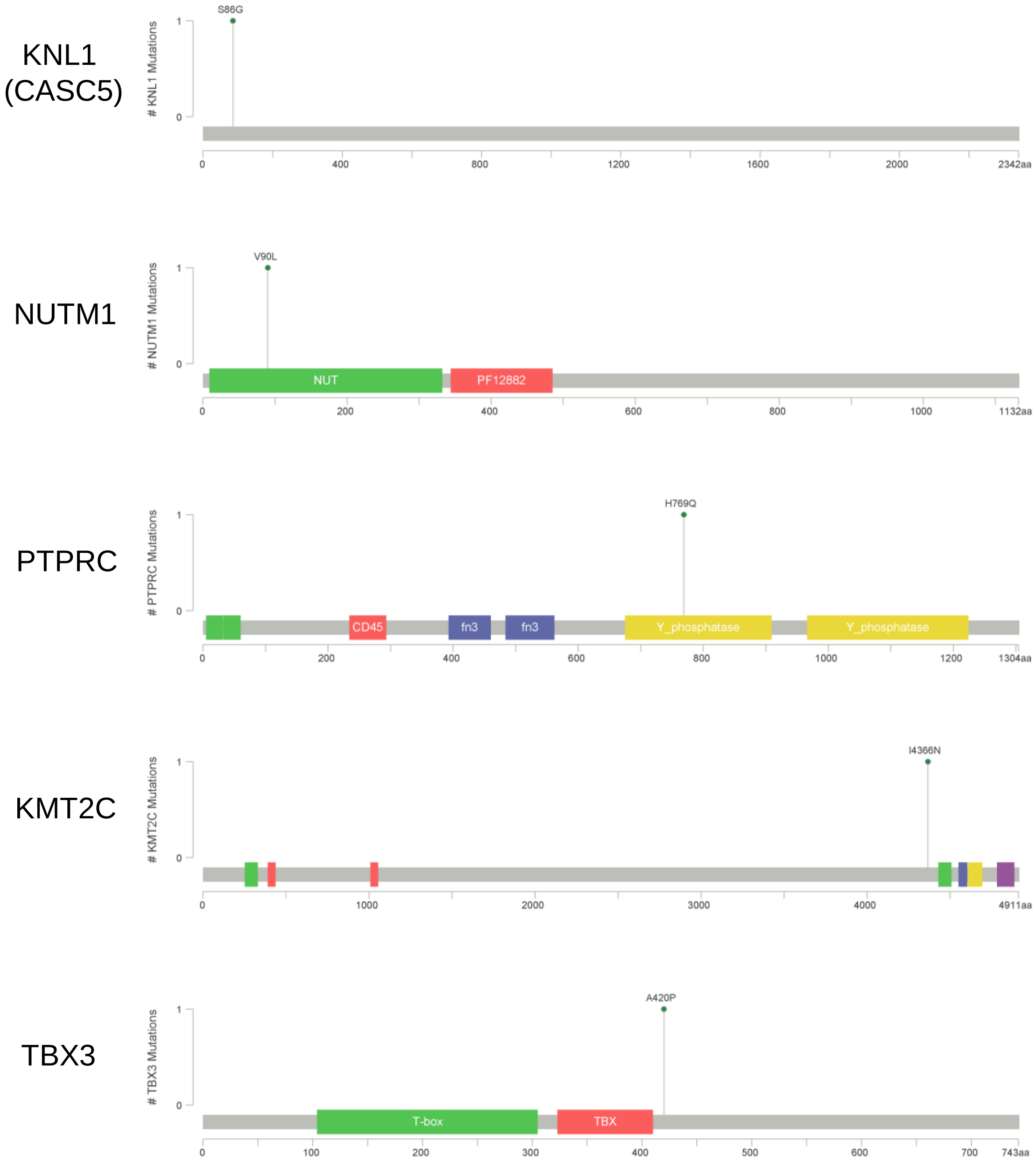


**Figure S2.** Mutation maps of the 5 driver genes of the prostate BCC.

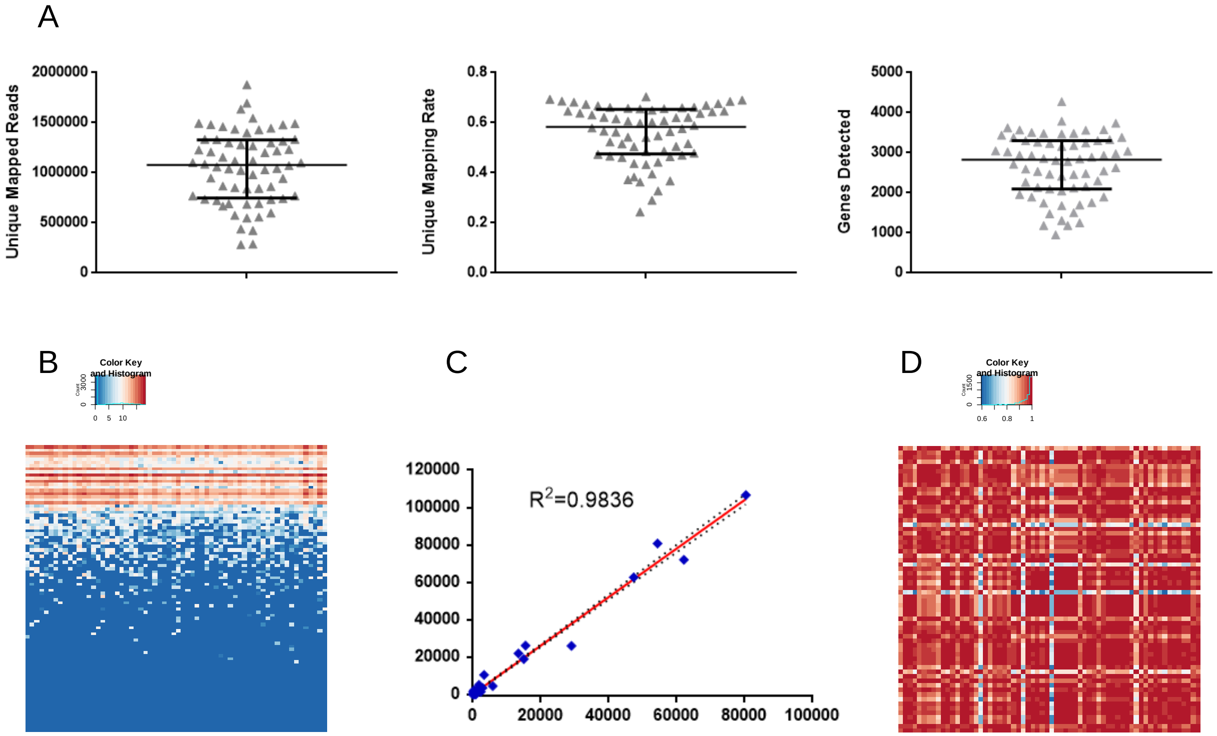


**Figure S3.** Quality assessment of single-cell RNA-Seq data of the prostate BCC. (A) Distribution of unique mapped reads, mapping rates, and the numbers of genes detected with FPKM > 1 in the single cells, with the median values and the first and third quartiles shown. (B) Heat-map showing the expression patterns of ERCC spike-ins, with each row represents a spike-in and each column represents a single cell. (C) An example of the correlation of the expression of ERCC spike-ins between two single cells. Dotted lines indicate the 95% confidence interval for the regression line. (D) Heat-map showing the Pearson correlation coefficients between single-cells based on ERCC spike-ins.

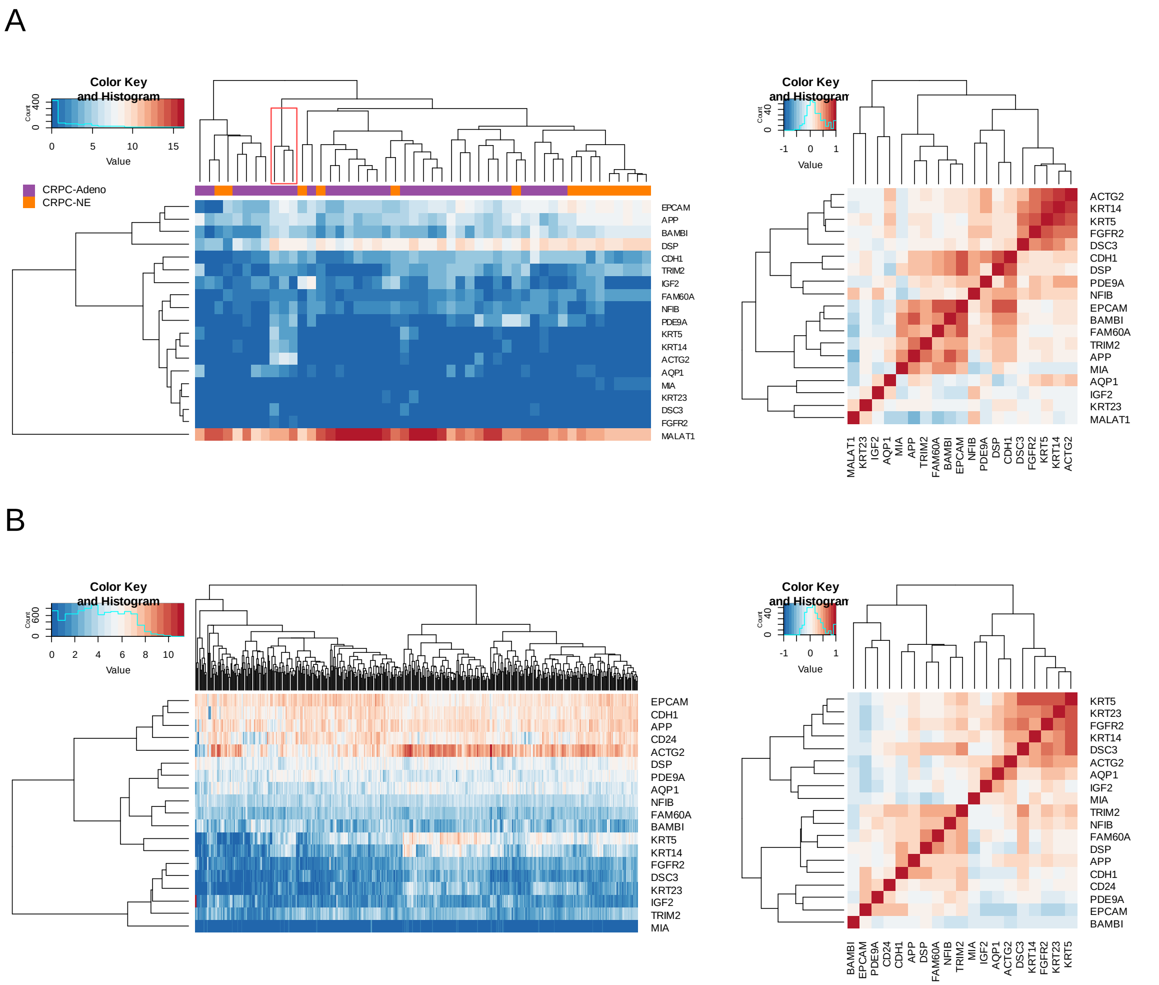


**Figure S4.** Gene expression (left) and correlation (right) patterns of BCC tumor specific genes in (A) castration-resistant prostate cancer (CRPC) and (B) TCGA PRAD samples. CRPC-Adeno: prostate adenocarcinomas; CRPC-NE: neuroendocrine prostate cancer. (Data from Beltran *et al*. Nat Med 2016; 22: 298-305 and TCGA Research Network)

A

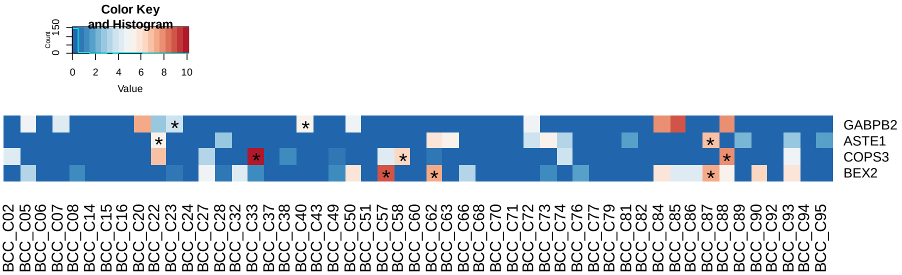


B

| Gene | AA change | Ref | Alt | Exome VAF | RNA-Seq Cell ID | Reference Count | Altered Count |
|---|---|---|---|---|---|---|---|
| *GABPB2* | M443T | T | C | 0.49 | BCC_C23 | 11 | 0 |
| | | | | | BCC_C40 | 0 | 11 |
| *ASTE1* | A57V | G | A | 0.40 | BCC_C22 | 18 | 0 |
| | | | | | BCC_C87 | 0 | 19 |
| *COPS3* | K105T | T | G | 0.45 | BCC_C33 | 0 | 115 |
| | | | | | BCC_C58 | 11 | 0 |
| | | | | | BCC_C88 | 19 | 0 |
| *BEX2* | D100N | C | T | 1.00 | BCC_C57 | 0 | 66 |
| | | | | | BCC_C62 | 0 | 19 |
| | | | | | BCC_C87 | 0 | 18 |

**Figure S5.** Detection of exonic mutations in single-cell RNA-Seq reads. (A) Gene expression heat-map of 4 genes where exonic variations are detected in the RNA-Seq reads. The stars indicate cells with exonic variations covered in RNA-Seq reads. (B) Summary of the exonic variations of prostate BCC in single-cell RNA-Seq data.
